# Impairment of a cyanobacterial glycosyltransferase that modifies a pilin results in biofilm development

**DOI:** 10.1101/2021.07.06.449578

**Authors:** Shiran Suban, Eleonora Sendersky, Susan S Golden, Rakefet Schwarz

**Affiliations:** The Mina and Everard Goodman Faculty of Life Sciences, Bar-Ilan University, Ramat-Gan, Israel, 5290002; Division of Biological Sciences, University of California, San Diego, La Jolla, CA 92093, USA; Center for Circadian Biology, University of California, San Diego, La Jolla, CA 92093, USA

**Keywords:** biofilm, glycosyltransferase, O-GlcNAc, PilA, pilus assembly, DNA competence, cyanobacteria, *Synechococcus elongatus* PCC 7942

## Abstract

A biofilm inhibiting mechanism operates in the cyanobacterium *Synechococcus elongatus*. Here, we demonstrate that the glycosyltransferase homolog, Ogt, participates in the inhibitory process – inactivation of *ogt* results in robust biofilm formation. Furthermore, a mutational approach shows requirement of the glycosyltransferase activity for biofilm inhibition. This enzyme is necessary for glycosylation of the pilus subunit and for adequate pilus formation. In contrast to wild-type culture in which most cells exhibit several pili, only 25% of the mutant cells are piliated, half of which possess a single pilus. In spite of this poor piliation, natural DNA competence was similar to that of wild-type, therefore, we propose that the unglycosylated pili facilitate DNA transformation. Additionally, conditioned medium from wild-type culture, which contains a biofilm inhibiting substance(s), only partially blocks biofilm development by the *ogt*-mutant. Thus, we suggest that inactivation of *ogt* affects multiple processes including production or secretion of the inhibitor as well as the ability to sense or respond to it.

**Originality-Significance Statement:** The molecular mechanisms that underlie biofilm development in cyanobacteria are just emerging. Using the cyanobacterium *S. elongatus* as a model, we demonstrate that glycosylation of the pilus subunit is crucial for the biofilm self-suppression mechanism, however, it is dispensable for DNA competence.

## Introduction

Post-translational modification is a pivotal mechanism that regulates protein function in all cell types, including the surface appendages of bacterial cells (Proft and Baker, 2009; Nothaft and Szymanski, 2010; Giltner et al., 2012). Surface modifications are likely to be important for cyanobacteria, which are photosynthetic prokaryotes that are highly prevalent in the environment, have an important role in global ecology (Garcia-Pichel et al., 2003; Falkowski et al., 2008; Braakman, 2019), and are found in microbial assemblages known as biofilms (Gorbushina, 2007; Stal et al., 2010; Mieszkin et al., 2013; Salta et al., 2013; Romeu et al., 2019; Kuhl et al., 2020). Among post-translational modifications, glycosylation is known to affect protein folding, localization and trafficking, protein solubility, antigenicity, biological activity and protein half-life (Blaser et al., 1986; Hounsell et al., 1996; Marceau and Nassif, 1999; Shental-Bechor and Levy, 2008; Vagin et al., 2009; Cummings, 2019).

Subunits of cell appendages required for motility are known to be glycosylated in diverse bacterial cells (Marceau and Nassif, 1999; Arora et al., 2005; Logan, 2006; Tytgat and Lebeer, 2014). For example, the flagellins FlaA and FlaB that comprise the flagellum of *Campylobacter jejuni* are heavily *N*-link glycosylated (Logan et al., 2002; Schirm et al., 2005). Glycosylation of *Campylobacter* flagellin is essential for flagellar assembly and consequent motility. Mutants defective in biosynthetic genes for the sugar moiety pseudominic acid in several strains of *Campylobacter* are non-motile and accumulate intracellular flagellin of reduced molecular mass due to lack of glycosylation (Hitchen et al., 2010). Another common glycosylation is addition of O-linked sugar onto pilus subunits; e.g., type IV pilins are O-glycosylated in some strains of *Pseudomonas aeruginosa, Neisseria gonorrhoeae* and *Neisseria meningitidis*, (DiGiandomenico et al., 2002; Power et al., 2003; Aas et al., 2007).

Studies of the molecular mechanisms underlying cyanobacterial biofilm development are currently emerging (Enomoto et al., 2014; Enomoto et al., 2015; Agostoni et al., 2016; Lacey and Binder, 2016; Enomoto et al., 2018). Our previous studies revealed a mechanism that represses biofilm formation in *Synechococcus elongatus* PCC 7942 (hereafter *S. elongatus*) (Schatz et al., 2013; Parnasa et al., 2016; Nagar et al., 2017; Sendersky et al., 2017; Parnasa et al., 2019). Inactivation of the gene Synpcc7942_2071 [recently designated as *pilB* (Yegorov et al., 2021)], which encodes an ATPase homologue of type II protein secretion (T2S) and/or type IV pilus assembly systems (T4P), impairs the inhibitory process, resulting in a mutant (PilB::Tn5) that develops robust biofilms (Schatz et al., 2013; Parnasa et al., 2016).

Factors that affect biofilm formation were recently shown to also affect multicellularity in *Synechocystis*. Motile strains of *Synechocystis* sp. PCC 6803 form floating aggregates termed flocs (Conradi et al., 2019) and impairment of PilB or the RNA chaperone homologue Hfq prohibited *Synechocystis* flocculation (Conradi et al., 2019). In contrast, inactivation of these components in *S. elongatus* elicited biofilm formation (Schatz et al., 2013; Yegorov et al., 2021), thus, these components impart different regulation of multicellularity in those cyanobacteria.

To identify additional components of the biofilm self-suppression mechanism of *S. elongatus*, a high throughput screen using a barcoded transposon library was employed (manuscript in preparation). This screen suggested involvement of several genes, including Synpcc7942_0051, in biofilm inhibition. Here, we demonstrate that the glycosyltransferase encoded by this gene is involved in glycosylation of the pilus subunit, PilA and discuss possible involvement of this enzyme in the biofilm inhibitory mechanism.

## Results

### Inactivation of Synpcc7942_0051 results in biofilm formation

A genetic screen using a barcoded transposon library suggested that inactivation of gene Synpcc7942_0051 causes biofilm development (manuscript in preparation). Briefly, this *S. elongatus* library of mutants was grown, a biofilm was allowed to form, and DNA extracted from the biofilm was analyzed by high throughput sequencing. Data analysis revealed mutants highly enriched in the biofilm, including a strain impaired in a gene encoding a glycosyltransferase homolog, O-linked β-N-acetylglucosamine (O-GlcNAc) transferase (Ogt). This enzyme is characterized by several tetratricopeptide (TPR) repeats at the N-terminus followed by two glycosyltransferase 41 domains [Fig. 1A, also see (Sokol and Olszewski, 2015)]. These domains are conserved in Ogt enzymes that catalyze attachment of O-GlcNAc to serine or threonine residues, a modification that is reversible due to an antagonistic activity by β-N-acetylglucosaminidases that remove O-GlcNAc (Golks and Guerini, 2008; Zeidan and Hart, 2010). However, the number of TPR repeats and the domain organization varies between enzymes [Fig. S1; (Lubas and Hanover, 2000; Iyer and Hart, 2003)].

**Figure 1:**
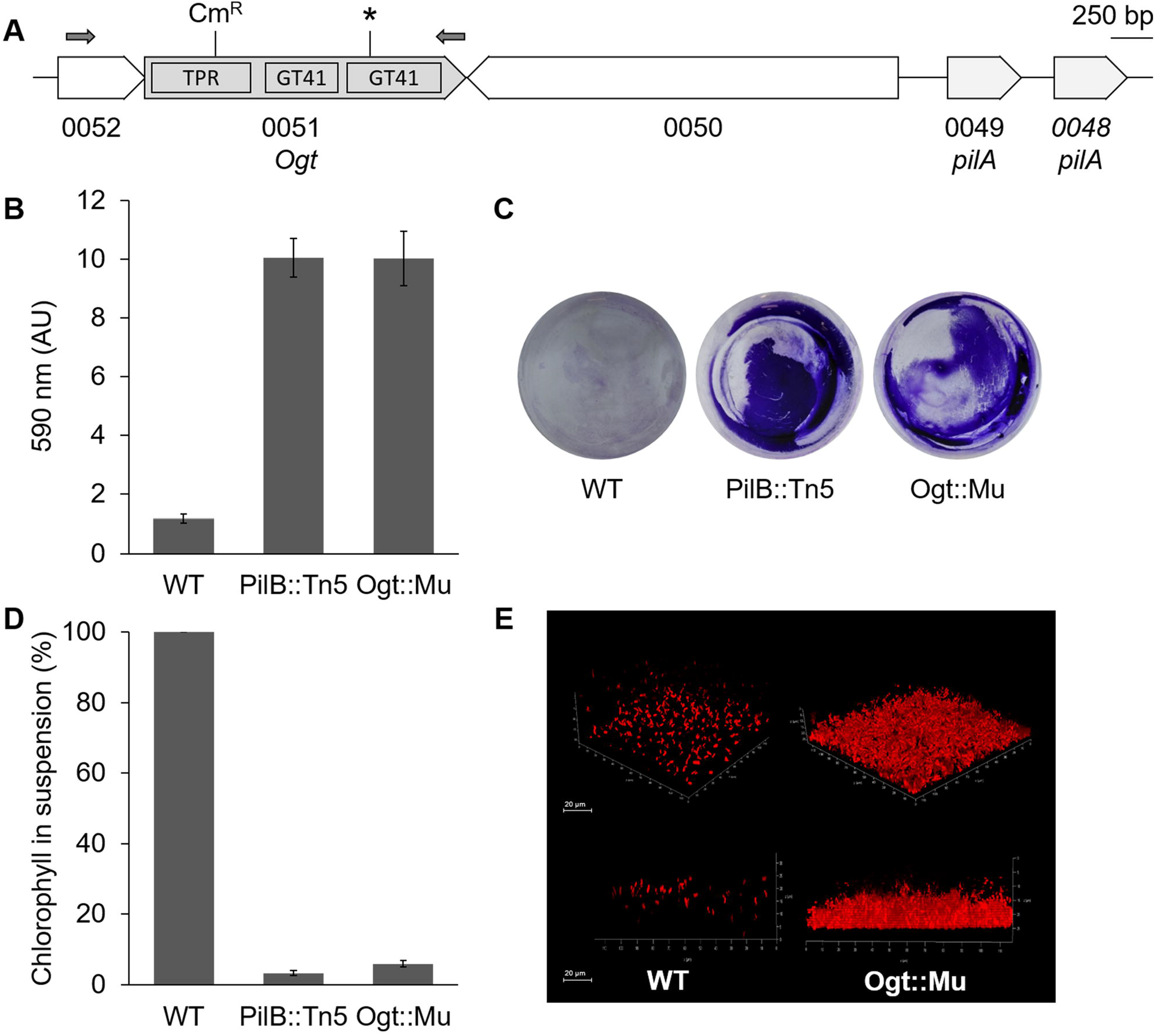
Inactivation of Synpcc7942_0051 results in biofilm formation. **A.** Genomic region of Synpcc7942_0051 (*ogt*). Tetratricopeptide (TPR) repeat region and conserved glycosyltransferase (GT41) domains are shown. The insertion point of a chloramphenicol resistance cassette (Cm^R^) is indicated. Asterisk indicates the position of the conserved lysine, K445. Small arrows indicate primers used to PCR amplify a DNA fragment for complementation (See Table S1). Genes Synpcc7942_0049/0048 encode the pilus subunit PilA. Synpcc7942_0050 and Synpcc7942_0052 are annotated ‘hypothetical’. **B.** Quantitation of biofilms using crystal violet staining. **C.** Biofilms formed at culture flask bottom as revealed by crystal violet staining. **D.** Percentage of chlorophyll in suspension as a proxy for biofilm development. Data in **B** and **D** represent average and standard error from 3 independent biological repetitions (with 3 technical repeats in each). **E.** Confocal microscopy imaging of biofilms formed in a flow cell. Red color represents auto-fluorescence (excitation: 588 nm; emission: 603-710 nm).

Enrichment of a mutant in the sessile subpopulation of the library implies that the particular strain is capable of biofilm development; however, it is possible that the mutant merely intercalated into and proliferated within a biofilm formed by other mutants in the library. Therefore, we insertionally inactivated Synpcc7942_0051 and tested biofilm formation by the mutant, Ogt::Mu, when grown in pure culture. The Ogt::Mu strain forms robust biofilms under static conditions (Fig. 1 B-D). Substantial biofilm development in flasks was revealed by crystal violet staining in Ogt::Mu similarly to PilB::Tn5 (Fig. 1 B&C). This assay suggested only trace cell adhesion or sugar-matrix deposition in wild-type (WT) culture (Fig. 1 B&C). Additionally, biofilms were quantified by assessment of the percentage of chlorophyll in suspension; about 95% of the total chlorophyll in cultures of this mutant is found in biofilm-cells, compared with 100% of the chlorophyll in WT cultures in planktonic cells (Fig. 1D). Total chlorophyll accumulated in the culture was not significantly different between WT and Ogt::Mu (Fig. S2).

Ogt::Mu did not reproducibly develop biofilms under the continuous bubbling conditions in which PilB::Tn5 biofilms are typically assayed (Schatz et al., 2013; Parnasa et al., 2016; Nagar et al., 2017; Sendersky et al., 2017; Parnasa et al., 2019). However, Ogt::Mu biofilms formed in a flow-cell system with continuous fresh BG-11 medium (Fig. 1E). In this setting, WT cells are observed in the groove of the flow-cell device but are not attached to the substratum (Fig. 1E, lower panel). Further characterization of the mutant was performed under static conditions.

Previously, we reported that conditioned medium from WT culture (hereafter CM) prohibited biofilm development by PilB::Tn5 under continuous bubbling, indicating that this mutant is capable of sensing and responding to biofilm-inhibitor(s) produced and secreted by the WT strain (Schatz et al., 2013; Parnasa et al., 2016). Here we tested the effect of CM on PilB::Tn5 and Ogt::Mu biofilm development under static conditions. In the absence of constant bubbling that maintained the cells suspended, PilB::Tn5 cells, which are non-piliated (Nagar et al., 2017), precipitated by gravity to the very bottom of the tube when grown in CM (Fig. 2, upper left tube). This pellet, however, was clearly distinct from the biofilm formed on the tube walls under fresh growth medium (Fig. 2, compare two upper tubes). In contrast to PilB::Tn5, the Ogt-mutant formed biofilms in CM, although not as robustly as under fresh medium and some cell pelleting was also observed in CM (Fig. 2, lower tubes). This observation suggests that Ogt::Mu is indifferent, to some extent, to the biofilm inhibitor present in CM from a WT culture.

**Figure 2:**
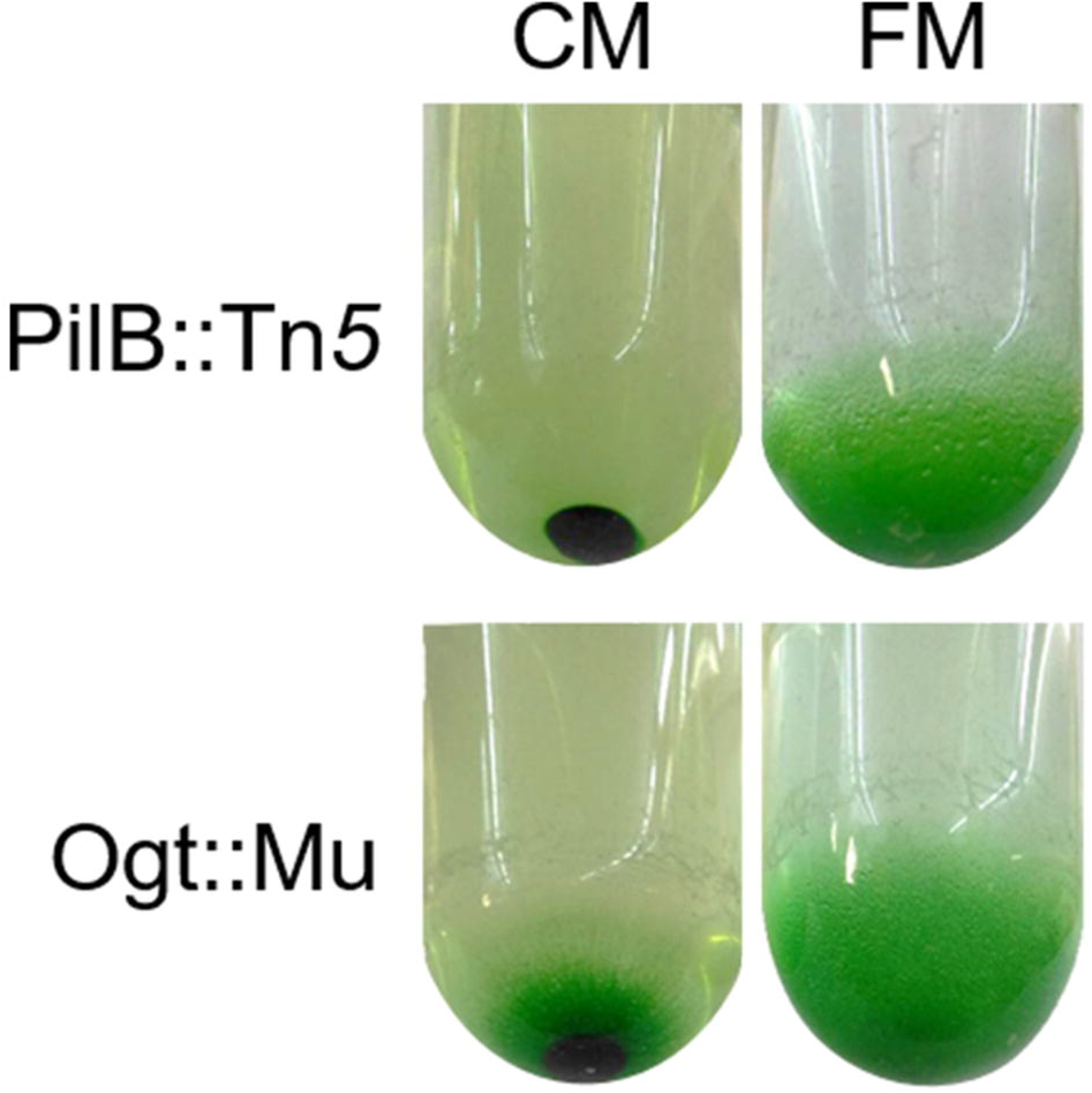
Biofilm development by PilB::Tn5 and Ogt::Mu under fresh medium (FM) and conditioned medium (CM) from WT culture. Cultures were imaged following 6 d of growth.

### Lysine 445 essential for catalysis is required for biofilm self-suppression

Biofilm development by Ogt::Mu indicates that Ogt takes part in the biofilm self-suppression mechanism that operates in *S. elongatus*. Next, we tested whether the predicted glycosyltransferase activity of Ogt is required for this process. We mutated the conserved lysine 445 in the C-terminal glycosyltransferase domain (Fig. 1A), which was previously demonstrated to be essential for catalytic activity (Clarke et al., 2008; Sokol and Olszewski, 2015). A gene encoding a lysine to alanine substitution (K445A) with its native regulatory region was introduced into the Ogt-mutant (Ogt::Mu/K445A, see Table S1). This strain formed biofilms similarly to Ogt::Mu, in contrast to the planktonic growth of strain Ogt::Mu/Ogt in which the native gene was inserted into the mutant (Fig. 3A). Because introduction *in trans* of *ogt* into Ogt::Mu restored planktonic growth, the biofilm-forming phenotype of Ogt::Mu is not due to effect of the antibiotic cassette inserted in *ogt* on downstream genes. Together, these data support requirement of Ogt glycosyltransferase activity for biofilm self-suppression.

**Figure 3:**
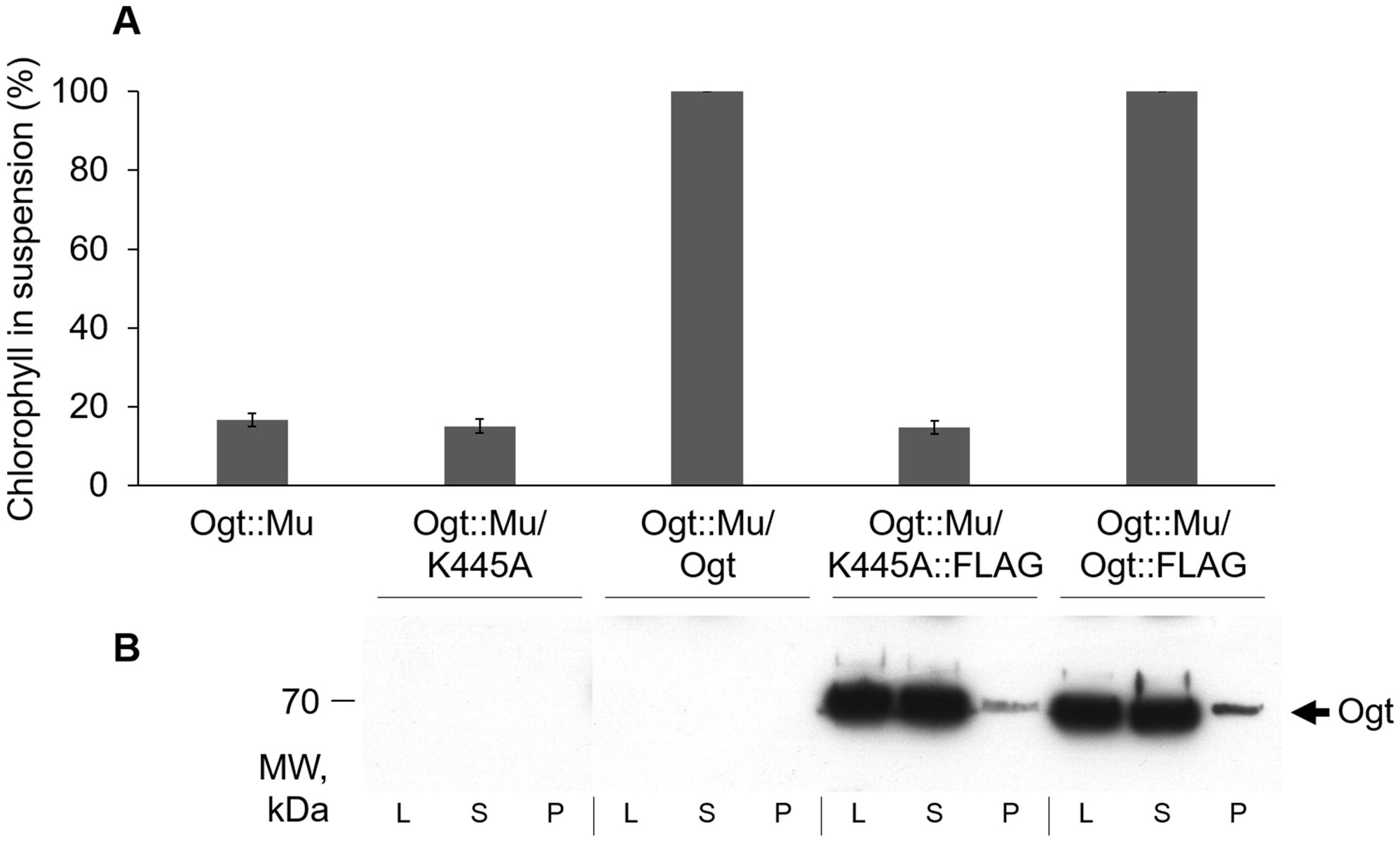
The glycosyltransferase activity of Ogt::Mu is required for biofilm self-suppression. **A.** Percentage of chlorophyll in suspension as a proxy for biofilm development. Data represent average and standard error from 3 independent biological repetitions (with 3 technical repeats in each). Strains analyzed: Ogt::Mu, Ogt-mutant harboring mutated and native Ogt (Ogt::Mu/ K445A and Ogt::Mu/Ogt, respectively) and corresponding strains with tagged mutated or native Ogt (Ogt::Mu/K445A::FLAG and Ogt::Mu/Ogt::FLAG, respectively). **B.** Western blot analysis of whole cell lysate (L), soluble (S) and pelletable (P) fractions using anti-FLAG antibodies. Arrow indicates the FALG-tagged Ogt protein (~70 kDa). Each lane corresponds to 2 μg chlorophyll.

Absence of restoration of planktonic growth in Ogt::Mu/K445A, could in principal result from lower stability or solubility of the mutated protein. Thus, we validated the presence of the mutant Ogt. A gene encoding C-terminal triple FLAG-tagged Ogt (hereafter Ogt::FLAG) was introduced with its native regulatory region into Ogt::Mu (Table S1). Western blot analysis using anti-FLAG antibodies indicated similar amount of tagged mutated and WT Ogt proteins, which were present mostly in the soluble cell fraction (Fig. 3B, Ogt::Mu/K445A::FLAG and Ogt::Mu/Ogt::FLAG, respectively). Moreover, the tagged-WT protein complemented the biofilm-forming phenotype of Ogt::Mu; in strain Ogt::Mu/Ogt::FLAG 100% of the chlorophyll is in planktonic cells (Fig. 3A). In contrast, strain Ogt::Mu/K445A::FLAG formed biofilms similarly to Ogt::Mu (Fig. 3A). Together, biofilm assays and Western analysis substantiate the requirement of the glycosyltransferase activity for the biofilm self-suppression process.

### Is Ogt involved in PilA glycosylation?

Our previous studies revealed involvement of the T4P assembly complex in the biofilm self-suppression mechanism operating in *S. elongatus*. Additionally, subunits of the pilus undergo glycosylation in diverse bacteria (Faridmoayer et al., 2007; Giltner et al., 2012; Harding et al., 2015; Elhenawy et al., 2016; Goncalves et al., 2018), and physical adjacency of genes encoding the pilus subunit and their cognate glycosyltransferases were reported e.g. in *Pseudomonas syringae* pv. tabaci 6605 (Nguyen et al., 2012) and in Ralstonia solanacearum (Elhenawy et al., 2016). Synteny analysis of 201 cyanobacterial genomes revealed physical adjacency of *ogt* and pilin genes in 77 (38%) of the genomes (see Fig. S3 for synteny in selected genomes). Given these facts, we tested whether Ogt is involved in PilA glycosylation in *S. elongatus*. Analysis of CM by gel-electrophoresis followed by silver staining indicated a major band of ~14 kDa in WT (Fig. 4A). This band, which is largely missing in PilB::Tn5 (Fig. 4A), corresponds to PilA (Nagar et al., 2017). In Ogt::Mu, however, a band of lower intensity and smaller MW compared to WT was detected (Fig. 4A). A similar shift as well as lack of glycosylation of the pilus subunit has been previously reported in diverse bacteria (Faridmoayer et al., 2007; Harding et al., 2015; Elhenawy et al., 2016) including cyanobacteria (Goncalves et al., 2018). The complemented strain Ogt::Mu/Ogt is characterized by a band similar in intensity and MW to that of WT. In contrast, a band akin to the one observed in Ogt::Mu was detected in Ogt::Mu/K445A (Fig. 4A). Taken together, these observations support involvement of Ogt in PilA glycosylation.

**Figure 4:**
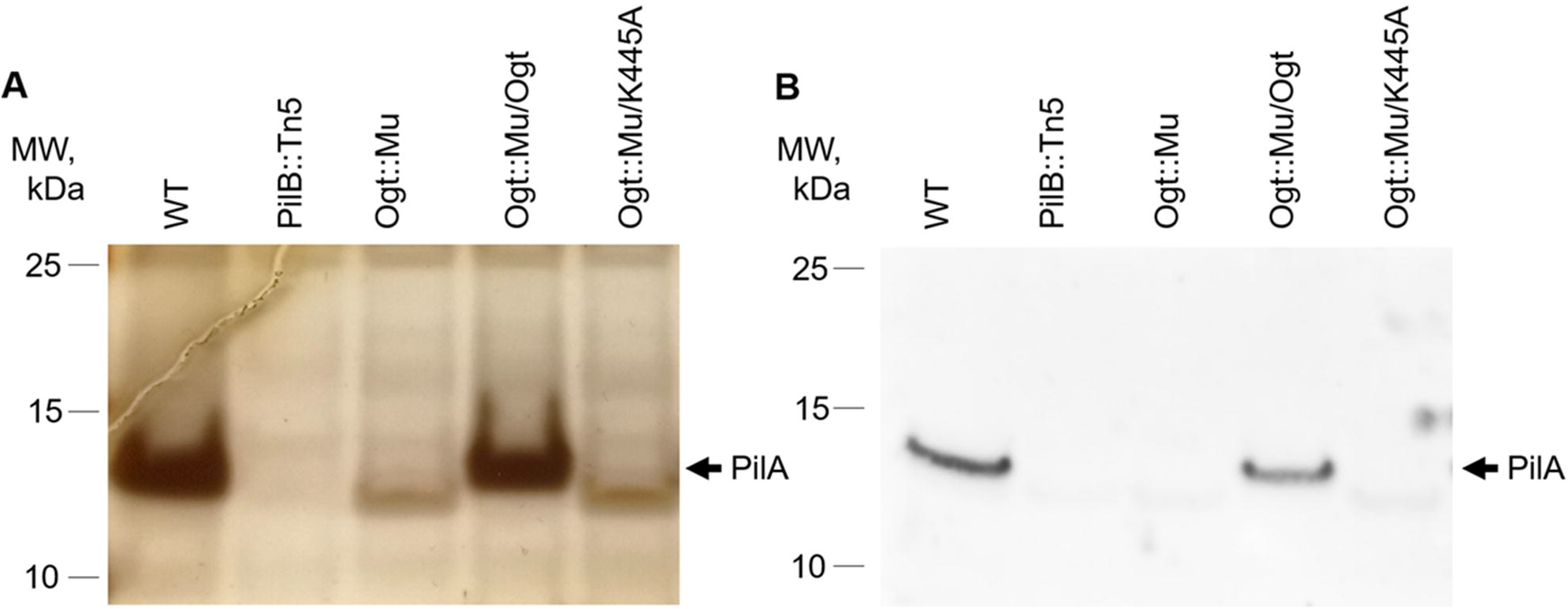
Ogt is involved in PilA glycosylation. Gel-electrophoresis of conditioned medium followed by silver staining **(A)** or GlcNAc detection **(B)**. Strains analyzed: *S. elongatus* (WT), *pilB*- and *ogt*-inactivated strains (PilB::Tn5 and Ogt::Mu, respectively) and Ogt::Mu into which native or mutated Ogt was introduced (Ogt::Mu/Ogt and Ogt::Mu/K445A, respectively). Each lane represents 0.5 ml of conditioned medium.

*In vitro* analysis has demonstrated that Ogt uses UDP-GlcNAc as a sugar donor (Sokol and Olszewski, 2015); therefore, we tested weather this glycosyltransferase is involved in addition of GlcNAc to PilA. Succinylated wheat Germ Agglutinin (succinyl-WGA), a lectin that detects GlcNAc, revealed a band of ~14 kDa in WT, supporting presence of GlcNAc residue(s) in PilA in this strain (Fig. 4B). This band is absent from Ogt::Mu; however, it was recovered in the *ogt* mutant bearing a native *ogt* gene (Fig 4B Ogt::Mu and Ogt::Mu/Ogt, respectively) but not with the active-site mutant allele (Fig. 4B, Ogt::Mu/K445A). In conclusion, the difference in MW (Fig. 4A) and GlcNAc detection (Fig. 4B) indicate involvement of Ogt in PilA glycosylation.

### Inactivation of ogt impairs pilus formation

Several studies demonstrated that subunits of cell appendages required for motility including T4P are glycosylated, and given the requirement of Ogt for PilA glycosylation we examined cell piliation in Ogt::Mu. TEM analyses of negatively stained whole cells revealed the presence of pili in WT [Fig. 5, (Nagar et al., 2017)]. The majority of the cells (96%) are piliated with most cells exhibiting several pili (see Table within Fig. 5). In contrast, only 25% of Ogt::Mu cells are piliated, half of which exhibit only a single pilus (Fig. 5). Introduction of *ogt* into the mutant largely restored pilus assembly (Fig. 5; 87% of the observed cells were piliated). Furthermore, in strain Ogt::Mu/K445A, which does not glycosylate PilA (Fig. 4), 21% of the cells were piliated, similar to Ogt::Mu (Fig. 5). Together, TEM analyses of the different strains support requirement of Ogt activity for proper pilus formation.

**Figure 5:**
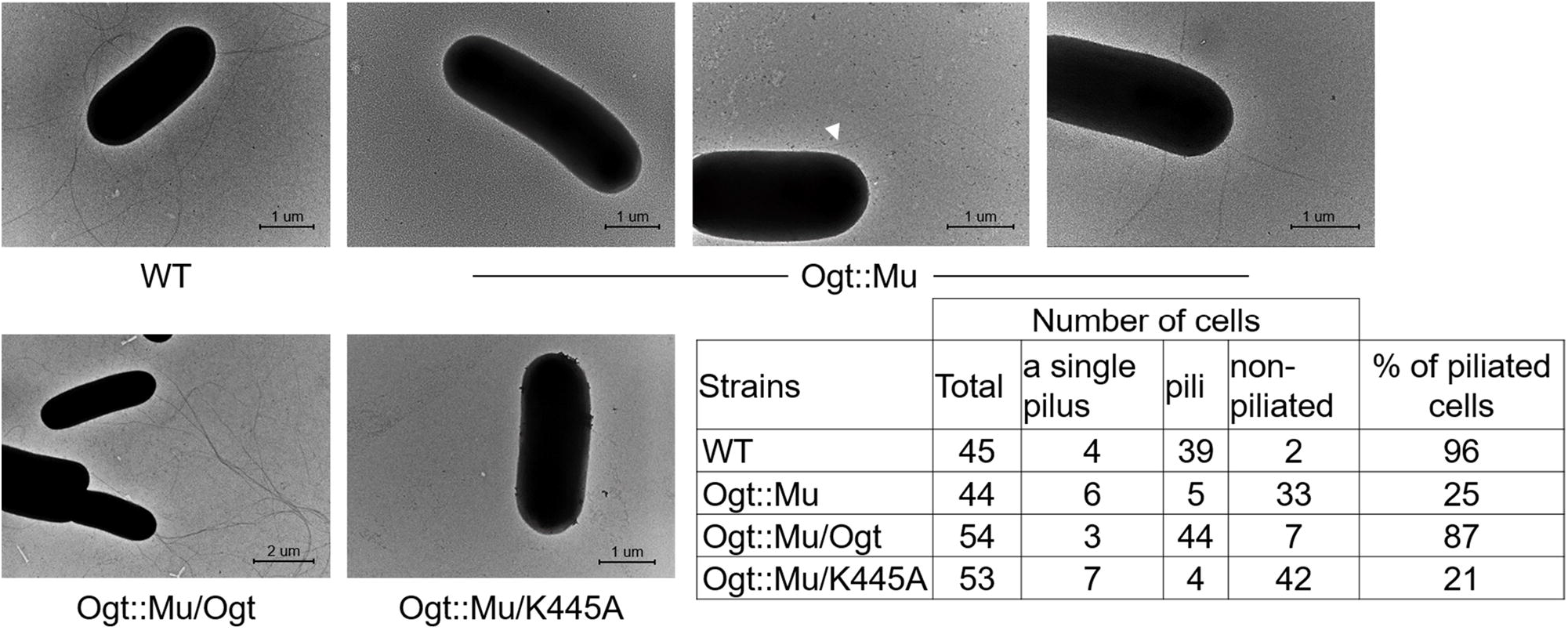
Ogt is required for adequate pilus assembly. TEM images of whole negatively stained cells and quantitative analysis of pili. Strains analyzed: *S. elongatus* (WT), *ogt*-inactivated strain (Ogt::Mu) and Ogt::Mu into which native or mutated *ogt* was introduced (Ogt::Mu/Ogt and Ogt::Mu/K445A, respectively). White arrow indicates the single pilus of an Ogt::Mu cell.

Type 4 pili are involved in DNA transformation (Ellison et al., 2018; Craig et al., 2019; Piepenbrink, 2019). As we previously reported (Nagar et al., 2017; Yegorov et al., 2021), non-plilated PilB::Tn5 completely lacks ability to take up external DNA (Fig. 6) in agreement with a recent study (Taton et al., 2020). Given the lower number of piliated cells and of pili per cell observed in Ogt::Mu compared to WT, we tested natural transformation competence of the mutant. Interestingly, despite the lower abundance of pili in Ogt::Mu, DNA competency of this strain is similar to WT (Fig. 6). Together, data suggest that the unglycosylated pili facilitate DNA transformation.

**Figure 6:**
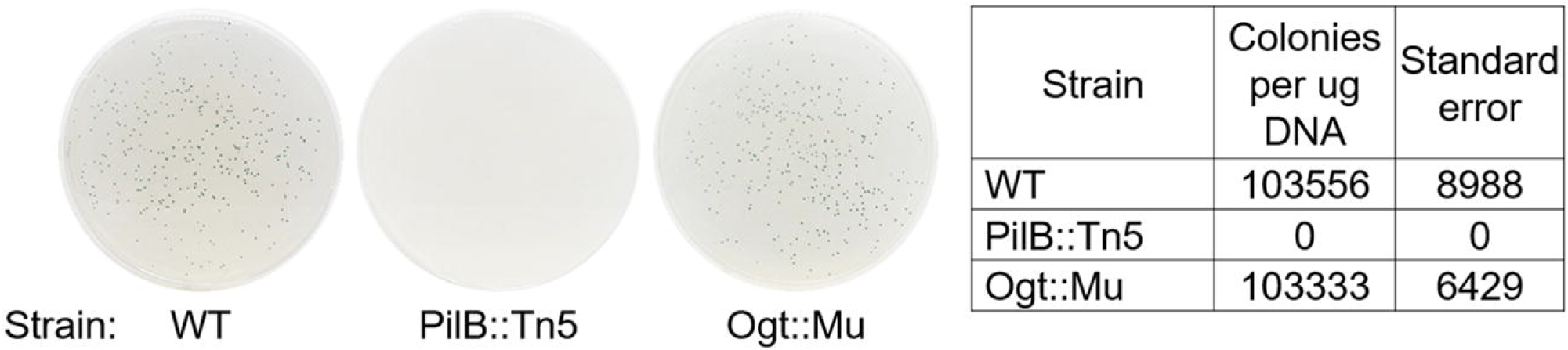
Inactivation of *ogt* does not affect natural DNA competence. DNA uptake by WT, PilB::Tn5 and Ogt::Mu. A shuttle vector capable of replication in *S. elongatus* was used, so that the transformation assay represents ability to take up the DNA without requirement for integration into the chromosome. WT and Ogt::Mu plates represent plating of 100 μL of 100^−2^ diluted transformation mixture whereas in the case of PilB::Tn5 200 μL of non-diluted mixture was plated. Data represent averages and standard error of 3 technical repeats.

Interestingly, inactivation of *ogt* affected fitness similarly to inactivation of genes encoding components of T4P and the DNA competence machinery (Table S2). The fitness data were collected using a randomly barcoded transposon-library that was subject to a large variety of growth conditions (Wetmore et al., 2015; Price et al., 2018). The abundance of a particular strain at the end of a growth experiment compared to its beginning served for fitness calculation. Genes are defined to have strong cofitness if they have similar fitness patterns (cofitness > 0.75). Very high cofitness values (0.89-0.99) were obtained for *ogt* and mutants in various *pil* genes encoding subunits of T4P assembly complex. Additionally, newly identified components of the T4P machinery that are essential for biofilm suppression [EbsA and Hfq, (Yegorov et al., 2021)] show high cofitness with Ogt (0.96 and 0.95, respectively, Table S2). These cofitness data support the involvement of Ogt in cellular processes related to the T4P assembly. Moreover, recently identified proteins required for natural transformation of *S. elongatus* [RntA and RntB, (Taton et al., 2020)] also show strong cofitness with Ogt (Table S2). Because inactivation of *ogt* did not affect DNA competence, an additional cellular function common with the Rnt proteins may be suggested.

## Discussion

### The glycosyltransferase Ogt is involved in biofilm self-suppression

The glycosyltransferase homologue Ogt is involved in the biofilm self-suppression mechanism operating in *S. elongatus* as manifested in biofilm development by the Ogt::Mu (Fig. 1). Sokol *et al*. (Sokol and Olszewski, 2015) reported that impairment of this glycosyltransferase results in cell settlement and formation of small cell aggregates, in contrast to development of robust biofilms reported here for the mutant. In both studies, cultures were analyzed under static conditions; however, other growth conditions (including temperature and light quality and intensity) varied between the studies. Additionally, we previously reported that freshly prepared iron stock is important for promotion of biofilm development, and we use it in our studies (Sendersky et al., 2017). Therefore, the conditions in the Sokol et al study may not have favored the biofilm development program beyond cell settling.

Previous studies indicated that impairment of pilus assembly is accompanied by low abundance of numerous proteins in the exo-proteome (Nagar et al., 2017; Yegorov et al., 2021). Furthermore, biofilm formation in non-piliated PilB::Tn5 was completely blocked by CM from a WT culture ((Schatz et al., 2013) and Fig. 2). Together, these observations support the hypothesis that the T4P complex in *S. elongatus* is engaged in protein secretion and is possibly involved in deposition of biofilm inhibitor(s) to the extracellular milieu. Distinct responses to CM, however, were observed for PilB::Tn5 and Ogt::Mu: Cell pelleting, apparently by gravity, was observed in PilB::Tn5 culture indicating complete inhibition of biofilm development, in contrast to the partial inhibition of biofilm formation in Ogt::Mu cultures that was manifested by some cell pelleting along with formation of biofilms on the tube walls (Fig. 2). These data may be explained by a pleiotropic effect of inactivation of *ogt* if Ogt affects the T4P apparatus as well as additional process(es) related to biofilm inhibition: e.g., the ability to properly respond to biofilm inhibitor(s) present in WT-CM. Pleiotropic effects resulting from abrogation of *ogt* in *S. elongatus* were reported by Sokol *et al*., (Sokol and Olszewski, 2015), including wider lumen of the photosynthetic membranes and high inorganic phosphate levels compared to WT. Our analysis revealed chlorophyll levels in Ogt::Mu similar to WT (Fig. S2), suggesting that biomass accumulation is not affected by impairment of the glycosyltransferase activity.

### Ogt is required for PilA glycosylation and adequate pilus formation

Several observations support involvement of *S. elongatus* Ogt in PilA glycosylation: PilA of Ogt::Mu is of lower molecular weight compared to WT (Fig. 4A) in agreement with lack of glycosylation in the mutant. Moreover, GlcNAc detection validated involvement of Ogt for PilA glycosylation (Fig. 4B). PilA glycosylation is commonly found in heterotrophic bacteria (Nothaft and Szymanski, 2010; Giltner et al., 2012). Recent studies in *Synechocystis* indicated that mutants defective in genes sll0141, sll0180 and slr0369, encoding homologues of TolC-related secretion systems, are impaired in PilA glycosylation. The particular role, however, of these proteins in the process of PilA glycosylation is unknown (Goncalves et al., 2018). An additional study of this cyanobacterium revealed that inactivation of sll0899 altered PilA glycosylation and impaired motility (Kim et al., 2009).

Impairment of PilA glycosylation (Fig. 4) abolished piliation in ~75% of the Ogt::Mu cells, whereas the rest were characterized by a single or few pili (Fig. 5). These observations show that despite the defective glycosylation, PilA secretion and assembly occurs albeit at low efficiency. Absence of motility of the lab strain of *S. elongatus* on agar plates prohibits assessment of the consequence of *ogt* inactivation on function of the assembled pili in motility.

The impact of PilA glycosylation on pilus assembly varies between different cyanobacteria. For example, deletion of sll0180 in *Synechocystis*, which results in lower MW PilA indicative of altered glycosylation, did not affect pilus formation as revealed by TEM (Goncalves et al., 2018). The deletion of *ogtA* in *Nostoc punctiforme* prevented accumulation of PilA, suggesting that PilA-glycosylation is required for its stability, and abolished cell motility. Additionally, transcription of *ogtA* of *N. punctiforme* is induced upon induction of development of motile filaments in this cyanobacterium, known as hormogonia. Together, these observations support requirement of OgtA for normal pilus function (Khayatan et al., 2015).

### Ogt is dispensable for DNA competence

Type IV pili are commonly involved in DNA uptake in diverse bacteria (Ellison et al., 2018; Craig et al., 2019; Piepenbrink, 2019) including cyanobacteria (Bhaya et al., 2000; Yoshihara et al., 2001; Schuergers and Wilde, 2015; Chen et al., 2020; Taton et al., 2020). For example, inactivation of *pilB*, completely eliminated pili and rendered the cells non-competent for DNA uptake (Nagar et al., 2017). A recent genome-wide screen of the same barcoded transposon library that identified *ogt* revealed the components required for natural transformation competence in *S. elongatus*, including various homologs of T4P assembly system (Taton et al., 2020). According to this screen, gene *ogt* is not required for DNA uptake, in agreement with the similar DNA competence of Ogt::Mu and WT reported here (Fig. 6). Though not completely abrogating pilus formation, inactivation of *ogt* severely reduced the number of piliated cells (Fig. 5). Therefore, a full suite of pili is not necessary to support maximal transformation competency. It is also possible that, as proposed for other organisms (Muschiol et al., 2015; Neuhaus et al., 2020), only particular pili serve in DNA uptake by *S. elongatus*, and *ogt* inactivation may not affect assembly of competence pili.

## Experimental Procedures

### Strains, culture conditions and biofilm quantification

Culture starters of *S. elongatus* PCC 7942 and all derived strains were bubbled with air enriched with CO_2_ as described previously (Sendersky et al., 2017). Starters were diluted with BG-11 to OD_750_ of 0.5 (30 ml culture in 100 ml flasks). Construction of mutants and details of molecular manipulations are provided in Table S1. Biofilms formed under static conditions at 28°C with incandescent light illumination (6 μmole photons*m^−2^ *s^−1^). Biofilms formed in standing cultures under the conditions indicated above and were quantified after 6 d growth as described (Parnasa et al., 2019), either by percentage of chlorophyll in suspension (Sendersky et al., 2017) or by crystal violet staining. For examination of biofilm formation in a flow cell system, initial cultures were diluted with BG-11 medium to OD_750_ of 1.0, and incubated for 48 h under 28°C and 6 μmole photons*m^−2^ *s^−1^. Subsequently, a constant flow of BG-11 at 5 ml/h was provided by a peristaltic pump. Biofilms were observed following 7 d from experiment initiation using Leica SP8 confocal microscope (excitation: 588nm; emission: 603-710 nm).

### Analysis of conditioned medium

Medium from a 50 ml culture was collected after centrifugation (5000 g, 10 min, room temperature). The supernatant was removed and passed through 0.22 mm filter, desiccated, and resuspended in 1.1 ml TE buffer supplemented with a protease inhibitor cocktail (Sigma, P8465-5ML). Analysis of conditioned medium by SDS-PAGE and silver staining was described previously (Schatz et al., 2013). Detection of *N*-Acetylglucosamine was performed as described (Cao et al., 2013) with the following modifications: 5% bovine serum albumin (BSA) in TBST was used as a blocking solution, 5 μg/ml biotinylated succinylated wheat germ agglutinin (Succinyl-WGA, VE-B-1025S, Vector Labs), which allows specific detection of GlcNAc was used rather than WGA, and horseradish peroxidase streptavidin (VESA-5014, Vector Labs) was adjusted to 3 μg/ml.

### Transmission electron microscopy

One day old cultures that had not yet initiated biofilm formation were sampled (10μl) and applied onto ultra-thin carbon coated grids. Following 5 minutes liquid was removed by blotting and cells were negatively stained twice with 2% fresh aquatic uranyl acetate (10 μl) for 1 min each step. Stain was removed by blotting, grids were briefly washed with double distilled water (10 μl) and left to dry over-night at room temperature. Images were acquired with a Technai G2 Fei transmission electron microscope, operating at 120 KV with a 1KX1K camera.

### DNA competence analysis

Assessment of DNA competence was performed essentially as described previously (Golden et al., 1987). Exponentially growing cells were centrifuged (5,000 × g for 8 min at room temperature), washed once with 10mM NaCl, and resuspended to an optical density at 750 nm (OD_750_) of 4.0. A shuttle vector (1,000 ng) was added to 600 μl of cells, which were gently agitated overnight at 28°C in the dark. Transformants were selected by plating on selective solid growth medium (50 μg/ml spectinomycin) supplemented with NaHCO3 (5 mM) and sodium thiosulfate (0.3%, wt/vol). The shuttle vector replicates autonomously, thus allowing the assessment of DNA uptake without a possible impact of the efficiency of DNA integration into the chromosome.

### Bioinformatics and confitness analysis

TPR repeats and glycosyltransferease domains were detected by TPRpred (Karpenahalli et al., 2007) and PFAM (Finn et al., 2014), respectively. Synteny analysis was performed using SyntTaX (Oberto, 2013). This tool is limited to analysis of 100 genomes at a time; therefore, analyses of 201 cyanobacterial genomes was performed in batches as follows: 50 genomes (grouped in SyntTaX under bacteria/cyanobacteria), and 100 and 51 (grouped in SyntTaX under bacteria:unclassified/cyanobacteira:unclassified). Cofitness data were recruited using the “Fitness Browser” web site (Wetmore et al., 2015; Price et al., 2018).

## Supporting information

Supplementary Information Suban et al

## Acknowledgments

Studies in the laboratories of Rakefet Schwarz and Susan Golden were supported by the program of the National Science Foundation and the U.S.-Israel Binational Science Foundation (NSF-BSF 2012823). This study was also supported by grants from the Israel Science Foundation (ISF 1406/14 and 2494/19) to Rakefet Schwarz.

## Legends to Supplementary Figures and Tables

**Figure S1: Domain organization of various Ogt enzymes.** TPR repeats (purple) and glycosyltransferase 41 domains (red) of cyanobacteria (*Synechococcus elongatus* PCC 7942, synpcc7942_0051; *Microcystis aeruginosa* PCC 7806, BH695_0839 and *Nostoc punctiforme* PCC 73102, Npun_F0677), gram positive bacteria (*Thermobaculum terrenum* strain ATCC BAA-798, Tter_2822 and *Listeria monocytogenes* strain EGD-e, lmo0688) and the gram-negative bacterium *Xanthomonas campestris* pv. campestris str. ATCC 33913, XCC0866.

**Figure S2: Inactivation of *ogt* does not affect biomass accumulation as measured by total chlorophyll.** Data represent average and standard error from 3 independent biological repetitions (with 3 technical repeats in each).

**Figure S3: Genomic analysis of different cyanobacteria indicates physical adjacency of *ogt* and *pilA* genes.** Selected genomic maps of the 201 cyanobacterial genomes analyzed by SynTax are presented.

**Table S1: Summary of cloning and mutational information**

* Insertional inactivation of Mu transposon, which confers Chloramphenicol resistance cassette (Cm^R^) was performed using the indicated vector from the unigene set (UGS) library (Holtman et al., 2005). Gene disruption in *S. elongatus* was obtained by transformation and replacement of the native gene by homologous recombination.

\# PCR products were cloned into the *Swa*I in NSII vector.

All cloning products were validated by PCR analyses and sequencing. Complete chromosome segregation of cyanobacterial strains were confirmed by PCR.

Lowercase letters in primers indicate the regions that encode the triple FLAG-tag.

**Table S2: Cofitness data of ogt-mutant.** Data were recruited from the Fitness Browser website (https://fit.genomics.lbl.gov/cgi-bin/myFrontPage.cgi) (Wetmore et al., 2015; Price et al., 2018).

## References

Aas, F.E., Vik, A., Vedde, J., Koomey, M., and Egge-Jacobsen, W. (2007) *Neisseria gonorrhoeae* O-linked pilin glycosylation: functional analyses define both the biosynthetic pathway and glycan structure. Mol Microbiol 65: 607–624.

Agostoni, M., Waters, C.M,. and Montgomery, B.L. (2016) Regulation of biofilm formation and cellular buoyancy through modulating intracellular cyclic di-GMP levels in engineered cyanobacteria. Biotechnol Bioeng 113: 311–319.

Arora, S.K., Neely, A.N., Blair, B., Lory, S., and Ramphal, R. (2005) Role of motility and flagellin glycosylation in the pathogenesis of *Pseudomonas aeruginosa* burn wound infections. Infect Immun 73: 4395–4398.

Bhaya, D., Bianco, N.R., Bryant, D., and Grossman, A. (2000) Type IV pilus biogenesis and motility in the cyanobacterium *Synechocystis* sp. PCC6803. Mol Microbiol 37: 941–951.

Blaser, M.J., Hopkins, J.A., Perez-Perez, G.I., Cody, H.J., and Newell, D.G. (1986) Antigenicity of *Campylobacter jejuni* flagella. Infect Immun 53: 47–52.

Braakman, R. (2019) Evolution of cellular metabolism and the rise of a globally productive biosphere. Free Radical Biology and Medicine 140: 172–187.

Cao, J., Guo, S., Arai, K., Lo, E.H., and Ning, M. (2013) Studying extracellular signaling utilizing a glycoproteomic approach : lectin blot surveys, a first and important step. Methods Mol Biol 1013: 227–233.

Chen, Z., Li, X., Tan, X., Zhang, Y., and Wang, B. (2020) Recent Advances in Biological Functions of Thick Pili in the Cyanobacterium *Synechocystis* sp. PCC 6803. Front Plant Sci 11: 241.

Clarke, A.J., Hurtado-Guerrero, R., Pathak, S., Schuttelkopf, A.W., Borodkin, V., Shepherd, S.M. et al. (2008) Structural insights into mechanism and specificity of O-GlcNAc transferase. EMBO Journal 27: 2780–2788.

Conradi, F.D., Zhou, R.Q,. Oeser, S., Schuergers, N., Wilde, A., and Mullineaux, C.W. (2019) Factors Controlling Floc Formation and Structure in the Cyanobacterium *Synechocystis* sp. Strain PCC 6803. J Bacteriol 201.

Craig, L., Forest, K.T., and Maier, B. (2019) Type IV pili: dynamics, biophysics and functional consequences. Nat Rev Microbiol 17: 429–440.

Cummings, R.D. (2019) “Stuck on sugars - how carbohydrates regulate cell adhesion, recognition, and signaling”. Glycoconjugate Journal 36: 241–257.

DiGiandomenico, A., Matewish, M.J., Bisaillon, A., Stehle, J.R., Lam, J.S., and Castric, P. (2002) Glycosylation of *Pseudomonas aeruginosa* 1244 pilin: glycan substrate specificity. Mol Microbiol 46: 519–530.

Elhenawy, W., Scott, N.E., Tondo, M.L., Orellano, E.G., Foster, L.J., and Feldman, M.F. (2016) Protein O-linked glycosylation in the plant pathogen *Ralstonia solanacearum*. Glycobiology 26: 301–311.

Ellison, C.K., Dalia, T.N., Vidal Ceballos, A., Wang, J.C., Biais, N., Brun, Y.V., and Dalia, A.B. (2018) Retraction of DNA-bound type IV competence pili initiates DNA uptake during natural transformation in *Vibrio cholerae*. Nat Microbiol 3: 773–780.

Enomoto, G., Okuda, Y., and Ikeuchi, M. (2018) Tlr1612 is the major repressor of cell aggregation in the light-color-dependent c-di-GMP signaling network of *Thermosynechococcus vulcanus*. Sci Rep 8: 5338.

Enomoto, G., Ni-Ni-Win, Narikawa, R., and Ikeuchi, M. (2015) Three cyanobacteriochromes work together to form a light color-sensitive input system for c-di-GMP signaling of cell aggregation. Proc Natl Acad Sci USA 112: 8082–8087.

Enomoto, G., Nomura, R., Shimada, T., Ni-Ni-Win, Narikawa, R., and Ikeuchi, M. (2014) Cyanobacteriochrome SesA Is a Diguanylate Cyclase That Induces Cell Aggregation in *Thermosynechococcus*. J Biol Chem 289: 24801–24809.

Falkowski, P.G., Fenchel, T., and Delong, E.F. (2008) The microbial engines that drive Earth’s biogeochemical cycles. Science 320: 1034–1039.

Faridmoayer, A., Fentabil, M.A,. Mills, D.C., Klassen, J.S., and Feldman, M.F. (2007) Functional characterization of bacterial oligosaccharyltransferases involved in O-linked protein glycosylation. J Bacteriol 189: 8088–8098.

Finn, R.D., Bateman, A., Clements, J., Coggill, P,. Eberhardt, R.Y., Eddy, S.R. et al. (2014) Pfam: the protein families database. Nucleic Acids Res 42: D222–230.

Garcia-Pichel, F., Belnap, J., Neuer, J., and Schanz, S. (2003) Estimates of global cyanobacterial biomass and its distribution. Algological Studies 109: 213–228.

Giltner, C.L., Nguyen, Y., and Burrows, L.L. (2012) Type IV pilin proteins: versatile molecular modules. Microbiol Mol Biol Rev 76: 740–772.

Golden, S.S., Brusslan, J., and Haselkorn, R. (1987) Genetic engineering of the cyanobacterial chromosome. Methods Enzymol 153: 215–231.

Golks, A., and Guerini, D. (2008) The O-linked N-acetylglucosamine modification in cellular signalling and the immune system. EMBO Reports 9: 748–753.

Goncalves, C.F., Pacheco, C.C., Tamagnini, P., and Oliveira, P. (2018) Identification of inner membrane translocase components of TolC-mediated secretion in the cyanobacterium *Synechocystis* sp. PCC 6803. Environ Microbiol 20: 2354–2369.

Gorbushina, A.A. (2007) Life on the rocks. Environ Microbiol 9: 1613–1631.

Harding, C.M., Nasr, M.A., Kinsella, R.L., Scott, N.E., Foster, L.J., Weber, B.S. et al. (2015) Acinetobacter strains carry two functional oligosaccharyltransferases, one devoted exclusively to type IV pilin, and the other one dedicated to O-glycosylation of multiple proteins. Mol Microbiol 96: 1023–1041.

Hitchen, P., Brzostek, J., Panico, M., Butler, J.A., Morris, H.R., Dell, A., and Linton, D. (2010) Modification of the *Campylobacter jejuni* flagellin glycan by the product of the Cj1295 homopolymeric-tract-containing gene. Microbiology 156: 1953–1962.

Holtman, C.K., Chen, Y., Sandoval, P., Gonzales, A., Nalty, M.S., Thomas, T.L. et al. (2005) High-throughput functional analysis of the *Synechococcus elongatus* PCC 7942 genome. DNA Res 12: 103–115.

Hounsell, E.F., Davies, M.J., and Renouf, D.V. (1996) O-linked protein glycosylation structure and function. Glycoconj J 13: 19–26.

Iyer, S.P.N., and Hart, G.W. (2003) Roles of the tetratricopeptide repeat domain in O-GlcNAc transferase targeting and protein substrate specificity. J Biol Chem 278: 24608–24616.

Karpenahalli, M.R., Lupas, A.N., and Soding, J. (2007) TPRpred: a tool for prediction of TPR-, PPR- and SEL1-like repeats from protein sequences. BMC Bioinformatics 8: 2.

Khayatan, B., Meeks, J.C., and Risser, D.D. (2015) Evidence that a modified type IV pilus-like system powers gliding motility and polysaccharide secretion in filamentous cyanobacteria. Mol Microbiol 98: 1021–1036.

Kim, Y.H., Kim, J.Y., Kim, S.Y., Lee, J.H., Lee, J.S., Chung, Y.H. et al. (2009) Alteration in the glycan pattern of pilin in a nonmotile mutant of *Synechocystis* sp. PCC 6803. Proteomics 9: 1075–1086.

Kuhl, M., Trampe, E., Mosshammer, M., Johnson, M., Larkum, A.W.D., Frigaard, N.U., and Koren, K. (2020) Substantial near-infrared radiation-driven photosynthesis of chlorophyll f-containing cyanobacteria in a natural habitat. eLife 9.

Lacey, R.F., and Binder, B.M. (2016) Ethylene Regulates the Physiology of the Cyanobacterium *Synechocystis* sp PCC 6803 via an Ethylene Receptor. Plant Physiology 171: 2798–2809.

Logan, S.M. (2006) Flagellar glycosylation - a new component of the motility repertoire? Microbiology 152: 1249–1262.

Logan, S.M., Kelly, J.F., Thibault, P., Ewing, C.P., and Guerry, P. (2002) Structural heterogeneity of carbohydrate modifications affects serospecificity of Campylobacter flagellins. Mol Microbiol 46: 587–597.

Lubas, W.A., and Hanover, J.A. (2000) Functional expression of O-linked GlcNAc transferase - Domain structure and substrate specificity. J Biol Chem 275: 10983–10988.

Marceau, M., and Nassif, X. (1999) Role of glycosylation at Ser63 in production of soluble pilin in pathogenic Neisseria. J Bacteriol 181: 656–661.

Mieszkin, S., Callow, M.E., and Callow, J.A. (2013) Interactions between microbial biofilms and marine fouling algae: a mini review. Biofouling 29: 1097–1113.

Muschiol, S., Balaban, M., Normark, S., and Henriques-Normark, B. (2015) Uptake of extracellular DNA: competence induced pili in natural transformation of Streptococcus pneumoniae. Bioessays 37: 426–435.

Nagar, E., Zilberman, S., Sendersky, E., Simkovsky, R., Shimoni, E., Gershtein, D. et al. (2017) Type 4 pili are dispensable for biofilm development in the cyanobacterium *Synechococcus elongatus*. Environ Microbiol 19: 2862–2872.

Neuhaus, A., Selvaraj, M., Salzer, R., Langer, J.D., Kruse, K., Kirchner, L. et al. (2020) Cryo-electron microscopy reveals two distinct type IV pili assembled by the same bacterium. Nat Commun 11: 2231.

Nguyen, L.C., Taguchi, F,. Tran, Q.M., Naito, K., Yamamoto, M., Ohnishi-Kameyama, M. et al. (2012) Type IV pilin is glycosylated in *Pseudomonas syringae pv. tabaci* 6605 and is required for surface motility and virulence. Molecular Plant Pathology 13: 764–774.

Nothaft, H., and Szymanski, C.M. (2010) Protein glycosylation in bacteria: sweeter than ever. Nat Rev Microbiol 8: 765–778.

Oberto, J. (2013) SyntTax: a web server linking synteny to prokaryotic taxonomy. BMC Bioinformatics 14: 4.

Parnasa, R., Sendersky, E., Simkovsky, R., Waldman Ben-Asher, H., Golden, S.S., and Schwarz, R. (2019) A microcin processing peptidase-like protein of the cyanobacterium *Synechococcus elongatus* is essential for secretion of biofilm-promoting proteins. Environ Microbiol Rep 11: 456–463.

Parnasa, R., Nagar, E., Sendersky, E., Reich, Z., Simkovsky, R., Golden, S., and Schwarz, R. (2016) Small secreted proteins enable biofilm development in the cyanobacterium *Synechococcus elongatus*. Sci Rep 6: 32209.

Piepenbrink, K.H. (2019) DNA Uptake by Type IV Filaments. Front Mol Biosci 6: 1.

Power, P.M., Roddam, L.F., Rutter, K., Fitzpatrick, S.Z., Srikhanta, Y.N., and Jennings, M.P. (2003) Genetic characterization of pilin glycosylation and phase variation in *Neisseria meningitidis*. Mol Microbiol 49: 833–847.

Price, M.N., Wetmore, K.M., Waters, R.J., Callaghan, M., Ray, J., Liu, H. et al. (2018) Mutant phenotypes for thousands of bacterial genes of unknown function. Nature 557: 503–509.

Proft, T., and Baker, E.N. (2009) Pili in Gram-negative and Gram-positive bacteria - structure, assembly and their role in disease. Cell Mol Life Sci 66: 613–635.

Romeu, M.J., Alves, P., Morais, J., Miranda, J.M., de Jong, E.D., Sjollema, J. et al. (2019) Biofilm formation behaviour of marine filamentous cyanobacterial strains in controlled hydrodynamic conditions. Env Microbiol 21: 4411–4424.

Salta, M., Wharton, J.A., Blache, Y., Stokes, K.R., and Briand, J.F. (2013) Marine biofilms on artificial surfaces: structure and dynamics. Env Microbiol 15: 2879–2893.

Schatz, D., Nagar, E,. Sendersky, E., Parnasa, R., Zilberman, S., Carmeli, S. et al. (2013) Self-suppression of biofilm formation in the cyanobacterium *Synechococcus elongatus*. Environ Microbiol 15: 1786–1794.

Schirm, M., Schoenhofen, I.C., Logan, S.M., Waldron, K.C., and Thibault, P. (2005) Identification of unusual bacterial glycosylation by tandem mass spectrometry analyses of intact proteins. Anal Chem 77: 7774–7782.

Schuergers, N., and Wilde, A. (2015) Appendages of the cyanobacterial cell. Life (Basel) 5: 700–715.

Sendersky, E., Simkovsky, R., Golden, S., S, and Schwarz, R. (2017) Quantification of Chlorophyll as a Proxy for Biofilm Formation in the Cyanobacterium *Synechococcus elongatus*. Bio-protocol 7: www.bio-protocol.org/e2406 DOI:2410.21769/BioProtoc.22406.

Shental-Bechor, D., and Levy, Y. (2008) Effect of glycosylation on protein folding: a close look at thermodynamic stabilization. Proc Natl Acad Sci U S A 105: 8256–8261.

Sokol, K.A., and Olszewski, N.E. (2015) The Putative Eukaryote-Like O-GlcNAc Transferase of the Cyanobacterium *Synechococcus elongatus* PCC 7942 Hydrolyzes UDP-GlcNAc and Is Involved in Multiple Cellular Processes. J Bacteriol 197: 354–361.

Stal, L.J., Severin, I., and Bolhuis, H. (2010) The ecology of nitrogen fixation in cyanobacterial mats. Recent Adv Phototrophic Prokaryotes: 31–45.

Taton, A., Erikson, C., Yang, Y., Rubin, B.E., Rifkin, S.A., Golden, J.W., and Golden, S.S. (2020) The circadian clock and darkness control natural competence in cyanobacteria. Nat Commun 11: 1688.

Tytgat, H.L.P., and Lebeer, S. (2014) The Sweet Tooth of Bacteria: Common Themes in Bacterial Glycoconjugates. Microbiol Mol Biol Rev 78: 372–417.

Vagin, O., Kraut, J.A., and Sachs, G. (2009) Role of N-glycosylation in trafficking of apical membrane proteins in epithelia. Am J Physiol Renal Physiol 296: F459–469.

Wetmore, K.M., Price, M.N., Waters, R.J., Lamson, J.S., He, J., Hoover, C.A. et al. (2015) Rapid quantification of mutant fitness in diverse bacteria by sequencing randomly bar-coded transposons. mBio 6: e00306–00315.

Yegorov, Y., Sendersky, E., Zilberman, S., Nagar, E., Waldman Ben-Asher, H., Shimoni, E. et al. (2021) A Cyanobacterial Component Required for Pilus Biogenesis Affects the Exoproteome. mBio 12.

Yoshihara, S., Geng, X., Okamoto, S., Yura, K., Murata, T., Go, M. et al. (2001) Mutational analysis of genes involved in pilus structure, motility and transformation competency in the unicellular motile cyanobacterium *Synechocystis* sp. PCC 6803. Plant Cell Physiol 42: 63–73.

Zeidan, Q., and Hart, G.W. (2010) The intersections between O-GlcNAcylation and phosphorylation: implications for multiple signaling pathways. J Cell Sci 123: 13–22.

