## Supplementary Information Suban et al for "Impairment of a cyanobacterial glycosyltransferase that modifies a pilin results in biofilm development"

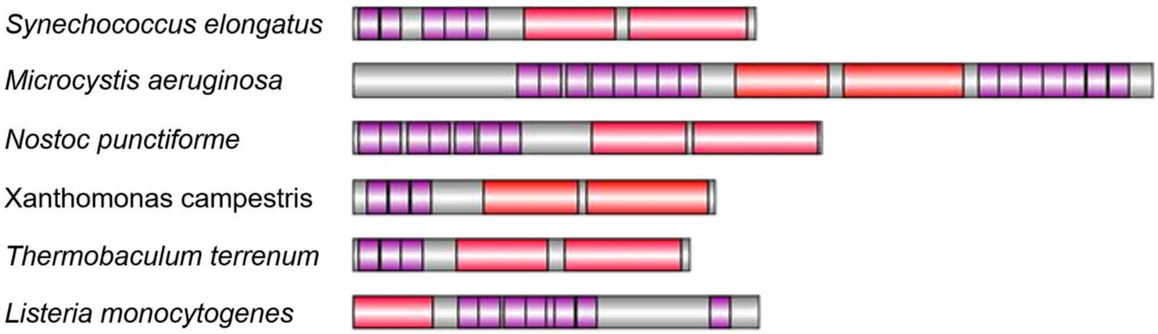

**Figure S1: Domain organization of various Ogt enzymes.** TPR repeats (purple) and glycosyltransferase 41 domains (red) of cyanobacteria (*Synechococcus elongatus* PCC 7942, synpcc7942_0051; *Microcystis aeruginosa* PCC 7806, BH695_0839 and *Nostoc punctiforme* PCC 73102, Npun_F0677), gram positive bacteria (*Thermobaculum terrenum* strain ATCC BAA-798, Tter_2822 and *Listeria monocytogenes* strain EGD-e, lmo0688) and the gram-negative bacterium *Xanthomonas campestris* pv. campestris str. ATCC 33913, XCC0866.

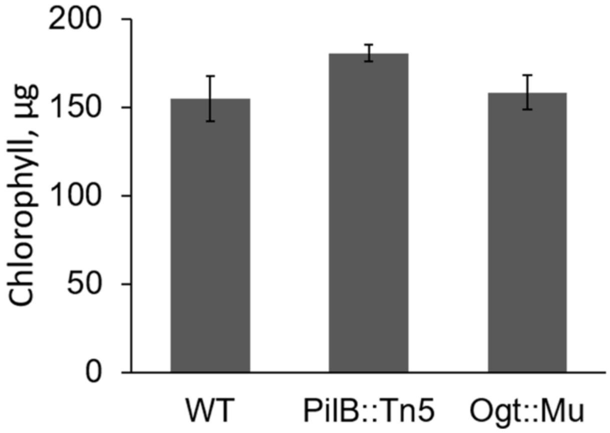

**Figure S2: Inactivation of *ogt* does not affect biomass accumulation as measured by total chlorophyll.** Data represent average and standard error from 3 independent biological repetitions (with 3 technical repeats in each).

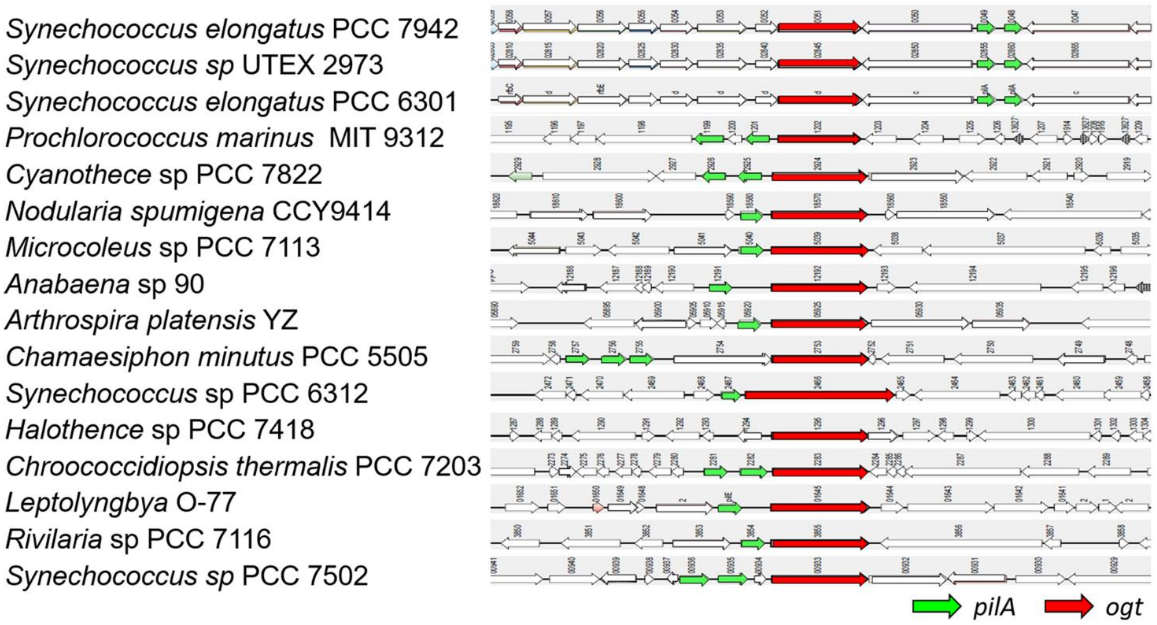

**Figure S3: Genomic analysis of different cyanobacteria indicates physical adjacency of *ogt* and *pilA* genes.** Selected genomic maps of the 201 cyanobacterial genomes analyzed by SynTax are presented.

| Gene disruption using transposon insertion vector | | |
|---|---|---|
| Inactivation of *ogt** (Synpcc7942_0051) | UGS vector 4E12 | Chloramphenicol resistance cassette ($Cm^R$) is inserted 649 bp downstream of the start codon |
| **Cloning of fragments encoding native and FLAG-tagged proteins** | | |
| | **Primer sequence (upper forward, lower reverse)** | **Aim** |
| *ogt* comp# | AGATTACATATTGGTGGAAGAG | PCR amplification |
| | TGCATGGTTAGAGAAGGTTTATC | |
| *ogt*::flag# | AGATTACATATTGGTGGAAGAG | PCR amplification |
| | TCActtatcgtcgtcatccttgtaatcgatatcgtgatccttgtaatccccatcgtgatccttgtaatcAAATCTAACTTCTTTATTGGCGCTC | |
| *ogt* K445A# | TAGTCGGCCAATAACCCAGGGATTTTGGAAGAGAAAAGCATCCGG | PCR amplification + Gibson assembly |
| | GAATAGTCACTAATTGCTCGACTTGCAACG | |
| *ogt* K445A::flag# | CGTTGCAAGTCGAGCAATTAGTGACTATTC | PCR amplification |
| | CTCCTGCCGGGGAGCTCCTTCATTTTCActtatcgtcgtcatccttgtaatcgatatcgtgatccttgtaatccccatcgtgatccttgtaatcAAATCTAACTTCTTTATTGGCGCTC | |
| **Additional primers** | | |
| *ogt* | AAGCCAGTTTGCAGATGACTCG | Examine segregation |
| | TTGAGGTGGTGGCGTGTAAGC | |
| *ogt* | TTATTGGTATGCACTGGCTG | Sequencing |
| | TTGAGGTGGTGGCGTGTAAGC | |
| NS 1 F/R | CGTCGAAGATGGAAAAGCTC | Examine segregation |
| | ATTGACCCGGTAGGGATTTC | |

**Table S1: Summary of cloning and mutational information**

* Insertional inactivation of Mu transposon, which confers Chloramphenicol resistance cassette ($Cm^R$) was performed using the indicated vector from the unigene set (UGS) library (Holtman et al., 2005). Gene disruption in *S. elongatus* was obtained by transformation and replacement of the native gene by homologous recombination.

### PCR products were cloned into the *Swa*I in NSII vector.

All cloning products were validated by PCR analyses and sequencing. Complete chromosome segregation of cyanobacterial strains were confirmed by PCR. Lowercase letters in primers indicate the regions that encode the triple FLAG-tag.

| # | Gene name synpcc7942_ | Name | Description | Cofitness |
|---|---|---|---|---|
| 1 | 1935 | *pilD* | type 4 prepilin peptidase 1. Aspartic peptidase. MEROPS family A24A | 0.97 |
| 2 | 1436 | | hypothetical protein | 0.96 |
| 3 | 0862 | *ebsA* | hypothetical protein | 0.96 |
| 4 | 2069 | *pilC* | fimbrial assembly protein PilC-like | 0.96 |
| 5 | 2486 | *rntA* | hypothetical protein | 0.96 |
| 6 | 2071 | *pilB1* | ATPase | 0.95 |
| 7 | 2479 | *pilA2* | pilin-like protein | 0.95 |
| 8 | 2451 | *pilO* | putative type IV pilus assembly protein PilO | 0.95 |
| 9 | 2484 | | hypothetical protein | 0.95 |
| 10 | 1139 | | hypothetical protein | 0.95 |
| 11 | 2452 | *pilN* | Tfp pilus assembly protein PilN-like | 0.95 |
| 12 | 0168 | *pilP* | hypothetical protein | 0.95 |
| 13 | 2453 | *pilM* | type IV pilus assembly protein PilM | 0.95 |
| 14 | 1926 | *hfq* | hypothetical protein | 0.95 |
| 15 | 2450 | *pilQ* | general secretion pathway protein D | 0.95 |
| 16 | 2485 | *rntB* | hypothetical protein | 0.95 |
| 17 | 1510 | *sigF1* | RNA polymerase sigma factor SigF | 0.94 |
| 18 | 1110 | | response regulator receiver domain protein (CheY-like) | 0.93 |
| 19 | 1525 | *typA* | GTP-binding protein TypA | 0.89 |
| 20 | 0049 | *pilA* | pilin polypeptide PilA-like | 0.89 |

**Table S2: Cofitness data of ogt-mutant.** Data were recruited from the Fitness Browser website (https://fit.genomics.lbl.gov/cgi-bin/myFrontPage.cgi) (Wetmore et al., 2015; Price et al., 2018).
